# A Latent Allocation Model for the Analysis of Microbial Composition and Disease

**DOI:** 10.1101/396903

**Authors:** Ko Abe, Masaaki Hirayama, Kinji Ohno, Teppei Shimamura

## Abstract

**Background:** Establishing the relationship between microbiota and specific disease is important but requires appropriate statistical methodology. A specialized feature of microbiome count data is the presence of a large number of zeros, which makes it difficult to analyze in case-control studies. Most existing approaches either add a small number called a pseudo-count or use probability models such as the multinomial and Dirichlet-multinomial distributions to explain the excess zero counts, which may produce unnecessary biases and impose a correlation structure taht is unsuitable for microbiome data.

**Results:** The purpose of this article is to develop a new probabilistic model, called BERMUDA (BERnoulli and MUltinomial Distribution-based latent Allocation), to address these problems. BERMUDA enables us to describe the differences in bacteria composition and a certain disease among samples. We also provide a simple and efficient learning procedure for the proposed model using an annealing EM algorithm.

**Conclusion:** We illustrate the performance of the proposed method both through both the simulation and real data analysis. BERMUDA is implemented with R and is available from GitHub (https://github.com/abikoushi/Bermuda).

## Background

Low-cost metagenomic and amplicon-based sequencing has provided a snapshots of microbial communities and their surrounding environments. One of the goals for case-control studies with microbiome data is to investigate whether cases differ from controls in the microbiome composition of a particular body ecosystems (e.g., the gut) and which taxa are responsible for any differences observed [1]. (Here, we use the generic term “taxa” to denote a particular phylogenetic classification.) These studies present microbiome data are represented as count data using operational taxonomic units (OTUs). The number of occurrences of each OTU is measured for each sample drawn from an ecosystem, and the resulting OTU counts are summarized at any level of the bacterial phylogeny, e.g., species, genes, family, order, etc. An important feature of these microbiome count data is that it is highly sparse—i.e., a very high proportion of the data entries are zero—which makes analyzing these data difficult.

A common strategy to handle these excess zeros is to add a small number called a pseudo-count [2]. Although adding a pseudo-count is simple and widely used, it can give the data an unnecessary bias to the data. Other strategies include modeling excess zeros using probability models [3, 4]. However, such models make an implicit assumption that all zeros can be explained by common probability models of microbial composition. Thus, such models cannot capture the important characteristic of individual differences in microbial composition.

### Contributions

This article propose a new probabilistic model, called BERnoulli and MUltinomial Distribution-based latent Allocation (BERMUDA), to address these problems. Our method can be regarded as a form of unsupervised learning. The contributions of our work are summarized below:

1. BERMUDA is a generative statistical model that allows a set of taxa to be explained by unobserved groups and can be used to find the inherent relationship between taxa and a specific disease and to generate microbiome count data through the model.
2. In BERMUDA, the abundance of each taxon can be viewed as a mixture of various groups, which enables us to describe the differences in bacteria composition between samples.
3. We provide a simple and efficient learning procedure for the proposed model using an annealing EM algorithm that reduces the local maxima problem inherent to the traditional EM algorithm. The software package that implements the proposed method in the R environment is available from GitHub (https://github.com/abikoushi/Bermuda).

We describe our proposed model and algorithm in “Methods” section. We also provide the efficiency of BERMUDA using synthetic and real data in “A Simulation Study” section and “Result for Real Data” section, respectively.

## Methods

### Proposed Model

Suppose that we observe a microbial count dataset with disease labels, {(*w*_*nk*_, *y*_*n*_); *n* = 1, *…, N, k* = 1, *…, K*)}, where *w*_*nk*_ is the abundance of the *k*-th taxon and *y*_*n*_ is a binary outcome such that *y*_*n*_ = 1 if the *n*-th sample has a certain disease and *y*_*n*_ = 0 otherwise. Let ***w***_*n*_ be the *k*-th row of matrix ***W*** = (*w*_*nk*_) and 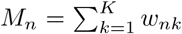 be the total reads count of the *n*-th sample.

We extract the associations between microbial composition and a specific disease by also supposing that there exist *L* latent clusters that vary with microbial composition and the disease risk. Let ***z**_n_* = (*z*_*n1*_, …, *z*_*nL*_)*^T^* be an indicator vector such that *z*_*nl*_ = 1 if the *n*-th sample is in the *l*-th class and *z*_*nl*_ = 0 otherwise. We then consider the following generative model:

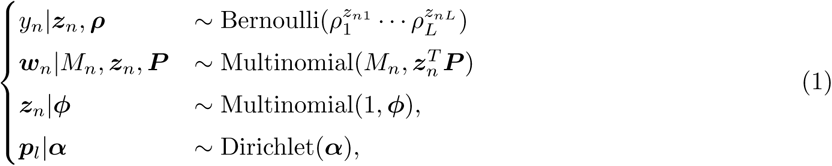

where ***ρ*** = (*ρ*_1_, *…, ρ_L_*)*^T^* is the probability of developing a certain disease, ***P*** = (*p*_*lk*_) (*l* = 1, *… L*) is an *L × K* matrix of the appearance probability of taxa, ***p***_*l*_ is the *l-*th row vector of matrix ***P***, ***ϕ*** = (*ϕ*_1_ *…, ϕ_L_*)*^T^* is a vector of each component’s mixing ratios, and ***α*** = (*α*_1_, *…, α_K_*)*^T^* is a vector of the hyperparameters of the Dirichlet prior distribution. Fig. 1 displays the plate notation for the proposed model. The gray node represents an observed variable and the white node represents an unobserved variable; the latent variable *z*_*n*_ affects both *y*_*n*_ and ***w***_*n*_.

**Figure 1:**
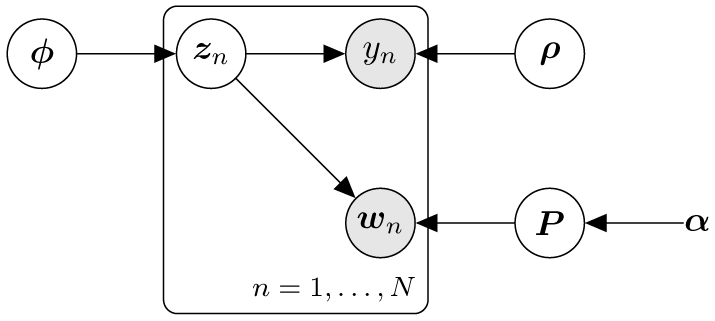
The plate notation for the proposed model

If the latent variable ***z***_*n*_ is given, the complete likelihood of this model is represented by the following formula:

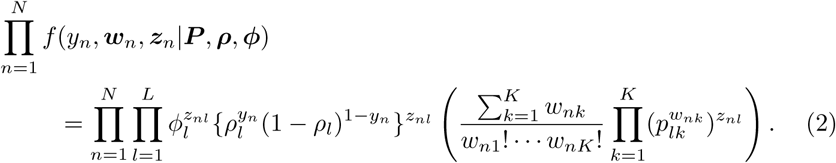

The posterior distribution is then proportional to:

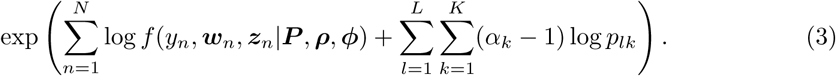

### Parameter Estimation

We find the maximum a posteriori probability (MAP) estimators, using an annealing EM (AEM) algorithm [5]. One advantage of using an AEM algorithm is that it reduces the local maxima problem from which the traditional EM algorithm suffers. In the E-step, using the inverse temperature 0 *< β ≤* 1, we calculate

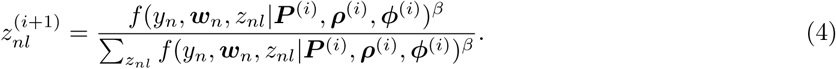

To simplify the explanation, we set *γ* = *α*_*k*_ - 1. From the logarithm of (3, in the M-step, we update the parameters using:

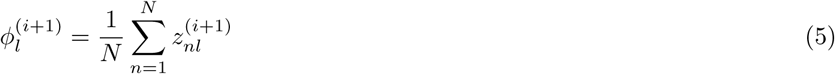

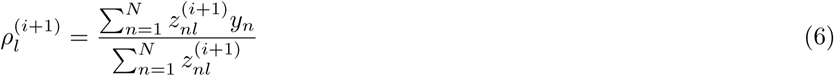

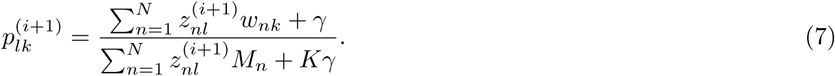

If *γ* = 0, MAP estimators are equivalent to MLEs.

A procedure of BERMUDA is then summarized as follows:

1. Set *β*.
2. Arbitrarily choose an initial estimate ***P***^(0)^, ***ϕ***^(0)^ and ***ρ***^(0)^. Set *i* ← 0
3. Iterate the following two steps until convergence: (a) E-step: Compute 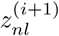 from (4). (b) M-step: Compute ***P***^(*i*+1)^, ***ϕ***^(*i*+1)^ and ***ρ***^(*i*+1)^ from (5), (6) and (7). Set *i ← i* = *i* + 1.
4. Increase *β*.
5. If *β <* 1, repeat from step 3; otherwise stop.

Let 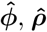 and 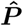 be MAP estimators of ***ϕ***, ***ρ*** and ***P***. If given ***w***_*n*_ and the estimators, we can evaluate the probability that the *n*-th sample has the target disease. The conditional probability is given by

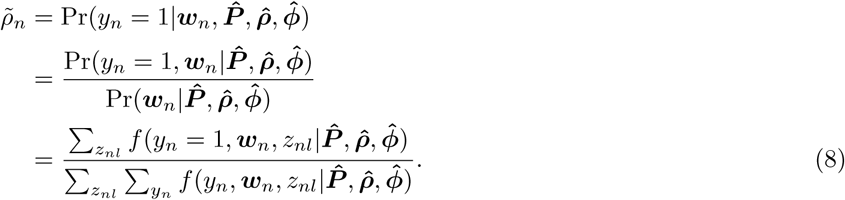

The advantage of using the Dirichlet prior distribution is that we can evaluate the abundance of the taxa whose abundance is exactly zero.

The *n*-th sample is then classified into the *l*-th class that maximizizes the conditional probability given by

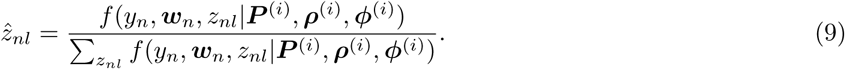

In fitting the model, it is important to choose an appropriate number for *L*. In this article, we use cross-validation to choose *L*. From (8), we can evaluate the probability that the *n*-th sample has the target disease. We can then evaluate the log-loss function represented by:

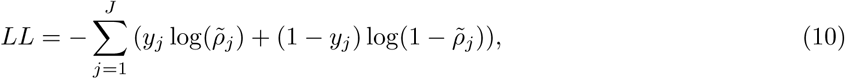

where *J* is an arbitrarily chosen subsample size for the validation data. We then select an *L* which minimizes (10) in this analysis.

### A Simulation Study

In this section, we generated synthetic data and evaluated the performance of our method in order to gain insights into the accuracy of the parameters estimated by the proposed method. A simulation study was conducted as follows. An i.i.d. sample is generated by (1) where we set *N* = 700, *M*_*n*_ = 10000, *L* = 7, *γ* = 10*^-9^*, ***ϕ*** = (1*/*7, *…,* 1*/*7)*^T^*, and ***ρ*** = (0, 3, 0.4, *…,* 0.9)*^T^*. ***P*** is chosen by a standard Dirichlet random number. We estimated the parameters from 10,000 replicates of the experiment.

Table 1 shows the mean and standard error (se) of the estimates for ***ρ*** and ***ϕ*** using the proposed method. It can be observed that the estimates are unbiased to the order of 1/100. Fig. 2 shows the relationship between estimates and true ***P*** in this simulation. In this figure, the points are arranged diagonally, which implies the estimator is unbiased. The overall accuracy of classification by *Ž*_*nl*_ (9) is 0.87. Thus, these results indicates that the proposed method can produce reasonable estimates and classify samples into true groups in this scenario.

**Table 1:**
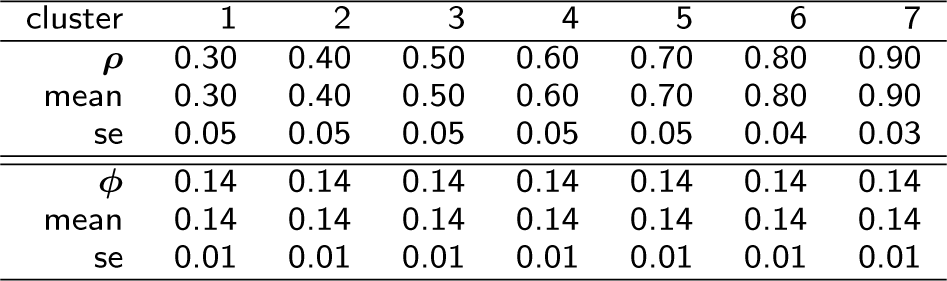
The mean and se *of* 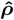 and 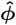

| cluster | 1 | 2 | 3 | 4 | 5 | 6 | 7 |
| --- | --- | --- | --- | --- | --- | --- | --- |
| $\rho$ | 0.30 | 0.40 | 0.50 | 0.60 | 0.70 | 0.80 | 0.90 |
| mean | 0.30 | 0.40 | 0.50 | 0.60 | 0.70 | 0.80 | 0.90 |
| se | 0.05 | 0.05 | 0.05 | 0.05 | 0.05 | 0.04 | 0.03 |
| $\phi$ | 0.14 | 0.14 | 0.14 | 0.14 | 0.14 | 0.14 | 0.14 |
| mean | 0.14 | 0.14 | 0.14 | 0.14 | 0.14 | 0.14 | 0.14 |
| se | 0.01 | 0.01 | 0.01 | 0.01 | 0.01 | 0.01 | 0.01 |

**Figure 2:**
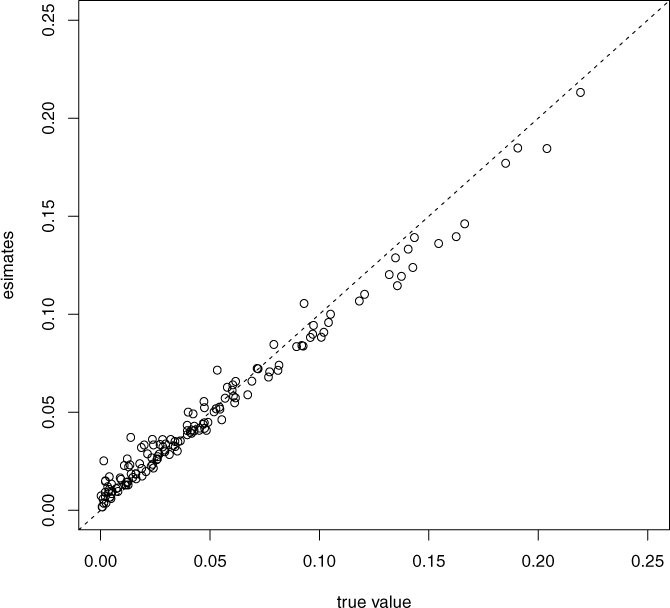
The comparison of true***P*** and mean of 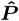

## Results for Real Data

We first seek to identify the gut dysbiosis in relation to development of Parkinson’s disease (PD), which is thought to be associated with intestinal microbiota. We analyzed intestinal microbial data in PD and controls in three different countries. Scheperjans *et al.* (2015) [6], Hill-Burns *et al.* (2017), [7] and Hopfner *et al.* (2017) [9] conducted case-control studies by sequencing the bacterial 16S ribosomal RNA gene in Finland, USA, and Germany, respectively.

The OTUs are then mapped to the SILVA taxonomic reference, version 132 (https://www.arb-silva.de/) and the abundances of genus-level taxa are calculated. We focused on the top 20 genera in terms of sample mean of normalized abundance *w*_*nk*_/*M*_*n*_ for 336 PD cases and 277 controls.

We set *γ* = 10*^-9^*, which is equivalent to giving a weakly informative prior. The number of components *L* = 6 is selected using 10-fold cross-validation (Fig 3). Fig. 3 shows the log-loss functions for different numbers of the components *L*.

**Figure 3:**
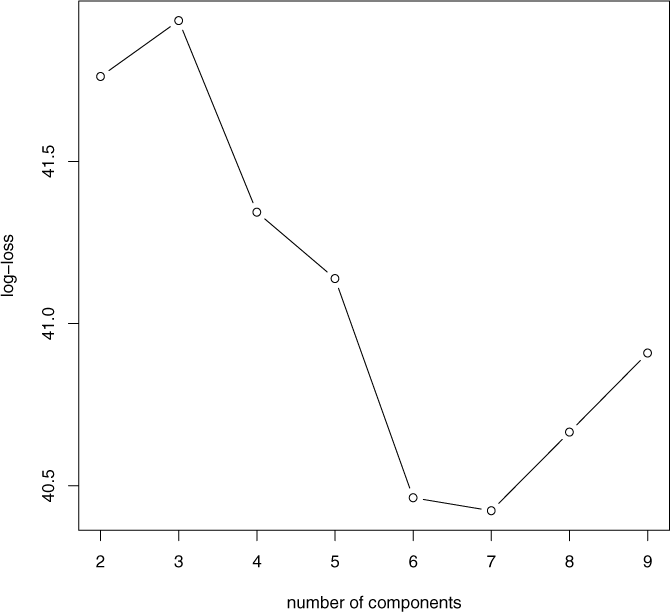
The behavior of the log-loss functions given by different numbers of components *L*

Fig. 4 presents the estimated appearance probabilities of the 20 genera. The clusters are sorted by estimated PD risk 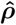 (Table 2). As displayed Fig. 4, the distribution of *Prevotella* is quite distinctive, being concentrated in the low-risk cluster of PD. *Faecalibacterium* also tends to be higher in the low-risk cluster. In contrast, *Akkermansia* is concentrated in the high-risk cluster.

**Table 2:**
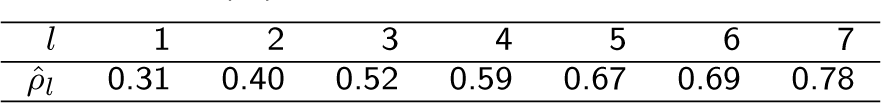
The estimated disease risk 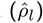 within each cluster

| $l$ | 1 | 2 | 3 | 4 | 5 | 6 | 7 |
| --- | --- | --- | --- | --- | --- | --- | --- |
| $\hat{\rho}_l$ | 0.31 | 0.40 | 0.52 | 0.59 | 0.67 | 0.69 | 0.78 |

**Figure 4:**
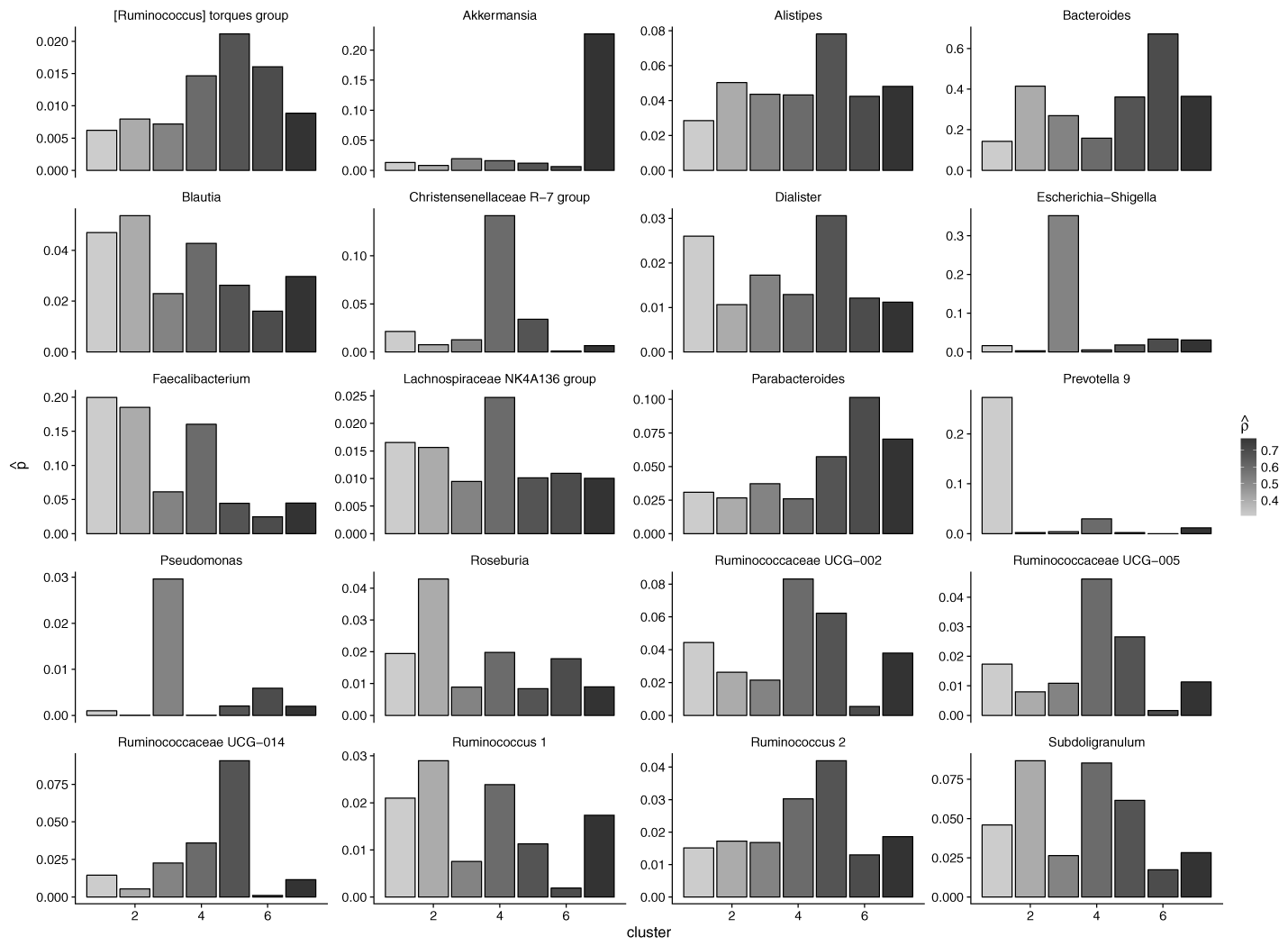
The appearance probability of the 20 genera 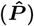

This result is consistent with the previous studies. Petrov *et al.* (2016) [10] reported that the gut microbiota of PD patients contained high levels of *Christensenella, Catabacter, Lactobacillus, Oscillospira*, and *Bifidobacteriumm*, and the control cluster was characterized by increased content of *Dorea, Bacteroides, Prevotella*, and *Faecalibacterium*. In family level analysis, Hill-Burns *et al.* (2017) [7] reported PD patients contained high levels of *Bifidobacteriaceae, Lactobacillaceae, Tissierellaceae, Christensenellaceae* and *Verrucomicrobiaceae* and low levels of *Lachnospiraceae, Pasteurellaceae*. *Scheperjans et al.*(2015) [6] reported PD patients contained high levels of *Lactobacillaceae, Verrucomicrobiaceae, Bradyrhizobiaceae* and *Ruminococcaceae* and low levels of *Prevotellaceae* and *Clostridiales Incertae Sedis IV*. *Akkermansia* belongs in *Verrucomicrobiaceae*. Of the *Verrucomicrobiaceae*, it has been suggested that *Akkarmansia* may be related to PD.

Thus, the analysis with real data demonstrate that the proposed method can identify the connection between the gut microbiota and the PD, with the results are strongly supported by the previous PD research.

## Conclusion

We proposed the new probabilistic model, called BERMUDA, for analyzing the relationship between microbiota and a specific disease. Although the existing approaches tend to underestimated individual differences in microbial composition, BERMUDA can take into account these differences and identify combinations of taxa rather than single taxa in the analysis of association with a specific disease risk. We demonstrated applicability of BERMUDA to microbial analyses with simulation and real data. The application of BERMUDA to gut microbiota data in PD and controls revealed that *Prevotella, Faecalibacterium*, and *Akkermansia* were associated with PD, which is consistent with previous studies. We expect that BERMUDA can be efficiently applieed to studies that seek for a causal association between gut dysbiosis and specific disease.

## Author’s contributions

KA and TS designed the proposed algorithm. KO and MH designed the experiments.

## Acknowledgements

This work was supported by Grants-in-Aid from the Ministry of Education, Culture, Sports, Science and Technology of Japan (MEXT); Ministry of Health, Labour and Welfare of Japan (MHLW); Japan Agency for Medical Research and Development (AMED), and the Hori Sciences and Arts Foundation.

## Availability of materials

BERMUDA is implemented with R and is available from GitHub (https://github.com/abikoushi/Bermuda).

## References

1. Brooks JP. Challenges for case-control studies with microbiome data. Annals of epidemiology, 2016; 26 (5): 336–341.

2. Xia F, Chen J.WK, Fung WK, Li H. A logistic normal multinomial regression model for microbiome compositional data analysi, Biometrics, 2013; 69 (4): 1053–63.

3. Chen EZ, Li H. A two-part mixed-effects model for analyzing longitudinal microbiome compositional data. Bioinformatics, 2016; 32 (17): 2611–7.

4. Paulson JN, Stine OC, Bravo HC, Pop M. Differential abundance analysis for microbial marker-gene surveys. Nat Methods, 2013; 10 (12): 1200–2.

5. Naonori Y, Nakano R. Deterministic annealing variant of the EM algorithm. Advances in neural information processing systems, 1995.

6. Scheperjans F, Aho V, Pereira PA, Koskinen K, Paulin L, Pekkonen E,… Kinnunen E. Gut microbiota are related to Parkinson’s disease and clinical phenotype. Movement Disorders, 2015; 30 (3): 350–358.

7. Hill-Burns EM, Debelius JW, Morton JT, Wissemann WT, Lewis MR, Wallen ZD,… Knight R. Parkinson’s disease and Parkinson’s disease medications have distinct signatures of the gut microbiome. Movement Disorders, 2017; 32 (5): 739–749.

8. Heintz-Buschart A, Pandey U, Wicke T, Sixel-Döring F, Janzen A, Sittig-Wiegand E,…, Wilmes P. The nasal and gut microbiome in Parkinson’s disease and idiopathic rapid eye movement sleep behavior disorder, Movement Disorders, 2018; 33 (1): 88–98.

9. Hopfner F, Künstner A, Müller SH, Künzel S, Zeuner KE, Margraf NG,…, Kuhlenbäumer G. Gut microbiota in Parkinson disease in a northern German cohort. Brain research, 2017; 1667, 41–45.

10. Petrov VA, Saltykova IV, Zhukova IA, Alifirova VM, Zhukova NG, Dorofeeva YB,…, Mironova YS. Analysis of Gut Microbiota in Patients with Parkinson’s Disease, Bulletin of experimental biology and medicine, 2016: 162(6): 734–737.

